# Accelerated growth of the scleractinian coral *Orbicella faveolata* under turbid and polluted water conditions

**DOI:** 10.1101/2024.11.20.624512

**Authors:** Luis Lizcano-Sandoval, Angela Marulanda-Gómez, Mateo López-Victoria, Alberto Rodriguez-Ramirez

**Affiliations:** Posgrado de Ciencias del Mar y Limnología, Universidad Nacional Autónoma de México, Puerto Morelos, Mexico; College of Marine Science, University of South Florida, St. Petersburg, FL, USA; Environmental Mapping Team, Spatial Informatics Group, Pleasanton, CA, USA; Department of Natural Sciences and Mathematics, Pontificia Universidad Javeriana Cali, Cali, Colombia; Institute of Biology and Chemistry of the Marine Environment (ICBM), Carl von Ossietzky University of Oldenburg, Oldenburg, Germany; Global Change Institute, The University of Queensland, St Lucia, QLD 4072, Australia; School of Biological Sciences, The University of Queensland, St Lucia, QLD 4072, Australia

**Keywords:** coral growth, luminescence, water quality, Varadero Reef, Cartagena Bay, Canal del Dique

## Abstract

Cartagena Bay, Colombia, experienced historical changes in water quality, mainly due to dredging and channelization of the Canal del Dique, which connects the bay with the Magdalena River. In 1981 the largest dredging events occurred in the Canal del Dique, which negatively affected the water quality in the bay. In this bay, the coral reef of Varadero depicts high coral diversity and large coral colonies, showing apparent signs of a healthy ecosystem, under highly turbid and polluted waters. Here, we evaluated the potential effects of historical water quality changes and environmental variables on the coral skeletal growth of *Orbicella faveolata*. We obtained coral growth information from 65 years in the period 1951–2015. Coral skeletal density was 0.73 g cm^-3^, linear extension 1.02 cm yr^-1^, and calcification 0.74 g cm^-2^ yr^-1^. After 1981, significant changes in coral growth and luminescence were found: skeletal density decreased (9.9%), linear extension increased (6.6%), and skeletal luminescence was less intense (1.6%). In overall, Atlantic Multidecadal Oscillation (AMO) data explained changes in skeletal density (8–10%) and linear extension (7–19%). Air temperature explained changes in skeletal luminescence (22–45%). Water flow of the Canal del Dique did not contribute significantly to any coral growth and luminescence variables. The growth of *O. faveolata* in Varadero Reef was characterized by low density skeletons and an accelerated linear extension, apparently influenced by the dredging events of 1981. Further research is required to understand coral growth responses and resilience to the conditions of high turbidity, pollution, and low light in Cartagena Bay.

## Introduction

Coral reefs are increasingly affected by a mixture of local and global stressors such as rising temperatures, disease outbreaks, eutrophication, acidification, and overexploitation that jeopardize their stability and persistence (Sully et al. 2019; Guan et al. 2020; Davis et al. 2021; Burke et al. 2023). In particular, water quality has been declining over time due to intensified human impacts on oceans and coastal areas (Halpern et al. 2019), which has exposed fringing coral reefs to higher levels of turbidity and pollution (Zweifler et al. 2021). While most reefs typically thrive in tropical and oligotrophic waters, ranging from shallow to mesophotic depths (Kahng et al. 2010; Guan et al. 2015), a fraction (12 %) of the world’s coral reefs are found in moderately turbid waters (0.080-0.127 Kd490) (Sully and van Woesik 2020). Further, evidence shows that some corals can survive under highly turbid and reduced light conditions (Camp et al. 2018; Burt et al. 2020; Zweifler et al. 2021) by finding refugia from thermal stress (Sully and van Woesik 2020), shifting from autotrophy to heterotrophy (Anthony and Fabricius 2000; Travaglione et al. 2023), reducing lipid content (Jones et al. 2020), photoacclimating to low light exposure (Jones et al. 2020; López-Londoño et al. 2021), and increasing their number of pigments per cell (Jones et al. 2020).

Despite these challenges, evidence shows that some coral reefs survive under highly turbid conditions (Camp et al. 2018; Burt et al. 2020; Zweifler et al. 2021). The coral reef facing the most extreme turbidity in the Caribbean is Varadero Reef, Colombia (Fig. 1) (López-Victoria et al. 2015; Zweifler et al. 2021). This well-developed reef exhibits typical signs of a healthy ecosystem, including high coral diversity, 45% coral cover, and large colonies of the coral *Orbicella faveolata* (López-Victoria et al. 2015; Pizarro et al. 2017). Varadero Reef exists in a light-limited environment, influenced by freshwater discharge and industrial wastewater flowing into Cartagena Bay (Fig. 1). Water quality in Cartagena Bay is highly variable both in seasonal and spatial scales (Tosic et al. 2019) (Fig. S1). Several parameters exceed threshold values that are not optimal for biotic health or sustaining life. Elevated concentrations of heavy metals, such as cadmium, lead, mercury, chromium, copper, nickel have been found in the water (Tosic et al. 2019), sediments (Cogua et al. 2012), and organisms (Alonso et al. 2000; Manjarrez et al. 2008). Cartagena Bay’s waters experience seasonal increases in nutrient levels and decreases in dissolved oxygen, occasionally reaching hypoxic conditions (Tuchkovenko and Lonin 2003; Tosic et al. 2019). Organic matter and high coliform counts have also been observed (Garay and Giraldo 1997; Tuchkovenko and Rendón 2002; Tosic et al. 2019). Domestic sewage and industrial wastewaters contribute pollutants to the bay, but it is the Canal del Dique that has caused significant declines in water quality (Tosic et al. 2018).

**Figure 1.**
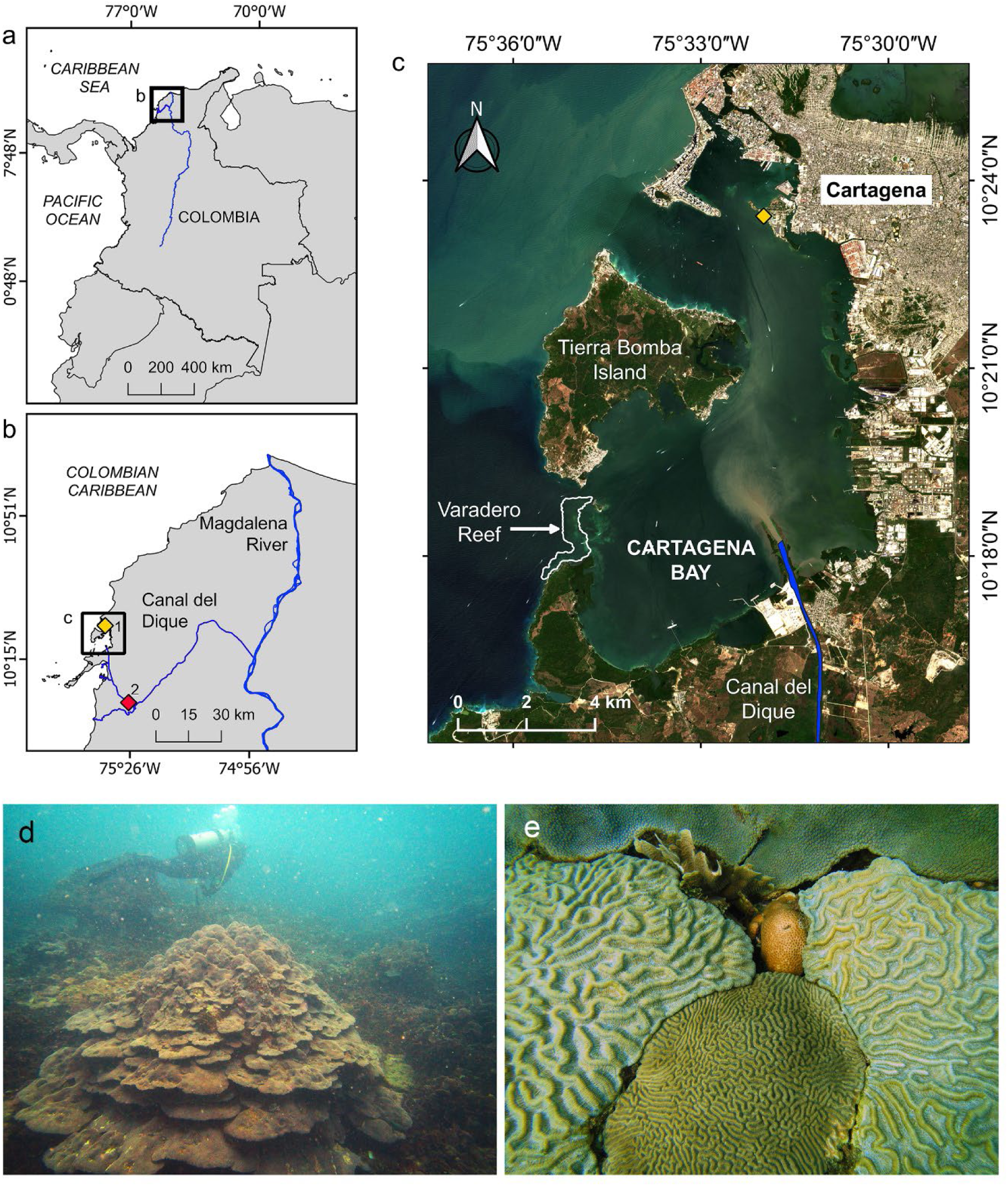
Geographic location of the Varadero coral reef in the Colombian Caribbean. (a-b) The connection between the Canal del Dique and the Magdalena River, and (c) location of Varadero reef within Cartagena Bay. Hydrological and meteorological stations are indicated by rhomboids: 1. Naval station (yellow), and 2. Santa Helena station (red). (d) *Orbicella faveolata* colonies growing in plate-like forms at 4 m depth with the presence of marine snow. (e) Corals competing for substrate in less than 1 m^2^: *Colpophyllia natans*, *Pseudodiploria clivosa*, *Agaricia tenuifolia*, *Siderastrea siderea* and *O. faveolata*.

The Canal del Dique connects the Magdalena River and Cartagena Bay since 1934, and it is a navigable channel of 115 km length (Mogollón 2002, 2013) (Fig. 1). It was fully channelized between 1950 and 1952, leading to the transformation of Cartagena Bay’s clear waters into turbid waters due to freshwater discharge (Mogollón 2013). Between 1981 and 1984, the Canal del Dique was rectified and expanded in both depth and width. During this period, nearly 47% of the soil and sediments dredged throughout the 20^th^ century were removed (Mogollón 2013). Annual maintenance is required to remove accumulated sediments, a practice that continues today. Between 1984 and 2010, the Canal del Dique increased water discharge by 28% and sediment load by 48%, contributing approximately 52 Mt of sediment to Cartagena Bay (Restrepo et al. 2018). While sedimentary evidence suggests that Cartagena Bay had a significant presence of scleractinian corals and seagrasses during the late Holocene (Martínez et al. 2010), recent changes in water quality have adversely affected these ecosystems. Seagrass cover in the bay declined by about 92% between 1935 and 2001 (Díaz and Gómez 2003), with no recent data available. Coral reefs here were once thought to be nonexistent, until recent studies (López-Victoria et al. 2015).

As turbid coral communities have survived under marginal conditions and shown resilience to climate change impacts (Zweifler et al. 2021), their corals may preserve critical information to support coral reef conservation strategies. Thus, a better understanding of the responses of such corals to recent changes in the environment and climate is pivotal to preserving the existence of these turbid reefs and managing the future resilience for other coral reef environments. Evaluating the resistance and recovery of coral reefs over disturbances and local environmental conditions is critical for developing effective conservation strategies (McClanahan et al. 2012). Massive corals respond to environmental changes by modulating their calcification rates and skeletal microstructure (e.g, exothecal dissepiments thickness), recording these variations in their skeleton (Barnes and Lough 1993; Carricart-Ganivet 2004; Dávalos-Dehullu et al. 2008). In corals of the genus *Orbicella*, seasonal temperature variations are recorded as changes in skeletal density bands, with low-density bands corresponding to cooler seasons and high-density bands indicating warmer seasons (Hudson et al. 1976; Carricart-Ganivet 2011). Each pair of bands represent one year of coral growth. Changes in skeletal luminescence also serve as a proxy for humic acid and sediment concentrations in the water (Grove et al. 2010; Tanzil et al. 2016). The ability of coral skeletons to record environmental responses provides an opportunity to track coral growth and reconstruct potential past disturbances.

The purpose of this paper was to document growth responses from the coral *O. faveolata* in the turbidity reef of Varadero, Colombia (Fig. 1) (López-Victoria et al. 2015; Zweifler et al. 2021). In this study, we measured coral growth parameters, including skeletal density, linear extension, calcification rates, and skeletal luminescence, in *Orbicella faveolata* to assess the effects of environmental changes and perturbations in Cartagena Bay on coral growth. We specifically analyzed the impacts of the Canal del Dique dredging in 1981 to provide insights on how corals have responded to changing environmental conditions.

## Material and methods

### Study area

The Varadero coral reef is located within Cartagena Bay, Colombian Caribbean (Fig. 1). Covering an area of approximately 1 km^2^, the reef presents up to 80% live coral cover, dominated by massive colonies of *O. faveolata* that can exceed 3 m in diameter (López-Victoria et al. 2015; Pizarro et al. 2017). This is a reef with conditions for coral settlement and growth in a limited space, exhibiting high coral species richness and density (Fig. 1e).

This coral reef is influenced by the turbid and polluted waters from the Canal del Dique, an extension of the Magdalena River (Fig. 1d, Fig. S1). The region experiences climate patterns that influence water quality in the bay. During the rainy season (September to December), salinity decreases while suspended particles and nutrient concentrations increase. In the windy season (January to April), water temperature decreases, and organic matter levels are high. During the transitional season (May to August), water temperature rises, and dissolved oxygen decrease (Tosic et al. 2019). Air temperatures are cooler from October to April and warmer from May to September (Fig. S2).

### Coral growth parameters

A single core of 6 cm diameter was extracted along the growth axis from four massive *O. faveolata* colonies at 3–4 m depth by scuba diving using a pneumatic drill. Coral density was estimated using a CT-scan, but one of the cores was cut and density was only possible to be estimated using the X-ray method.

One coral core (VAR1) was sliced using a circular saw into several slabs of 6 mm thick (prior to using the CT-scan method for coral density estimation). One slabs was selected to be radiographed using a conventional X-ray machine, and coral density was estimated according to Carricart-Ganivet and Barnes (2007). Radiation effects on the radiograph were corrected following Duprey (2012). The remaining three coral cores (VAR2, VAR3, VAR4) were imaged using a General Electric LightSpeed VCT 64 CT-scan as described by (Lizcano-Sandoval et al. 2019). The skeletal density of all four coral specimens was calibrated using aragonite density standards obtained from the shell of the giant clam *Tridacna maxima* (Lizcano-Sandoval et al. 2019).

For scanned coral cores, the best planes for observing the density banding pattern were selected by inspecting multi-planar reconstructions. For each core, one sagittal and one coronal image were chosen, and a single transect was drawn along the coral growth axis in each image to extract optical density values in Hounsfield units (HU). The HU values from both transects were averaged for each core. These averaged HU values were then converted to skeletal density (Lizcano-Sandoval et al. 2019). The CT-scan images were visualized and analyzed using RadiAnt DICOM Viewer 4.0.

The annual skeletal density pattern (Fig. 2) for orbicellid corals is characterized by a high-density band formed during the warm season and a low-density band formed during the cold season (Carricart-Ganivet et al. 2000; Carricart-Ganivet 2004). Here, we matched the highest and lowest density peaks along the coral skeleton to late June and February, respectively, to be consistent with the climatology of Cartagena Bay (Fig. S2). For both CT-scan and X-ray images, coral years were counted retrospectively. Skeletal density (g cm^-3^), linear extension (cm yr^-1^), and calcification rates (g cm^-2^ yr^-1^) were estimated annually. Linear extension was calculated from early January to late December of each year, while calcification was determined as the product of skeletal density and linear extension.

**Figure 2.**
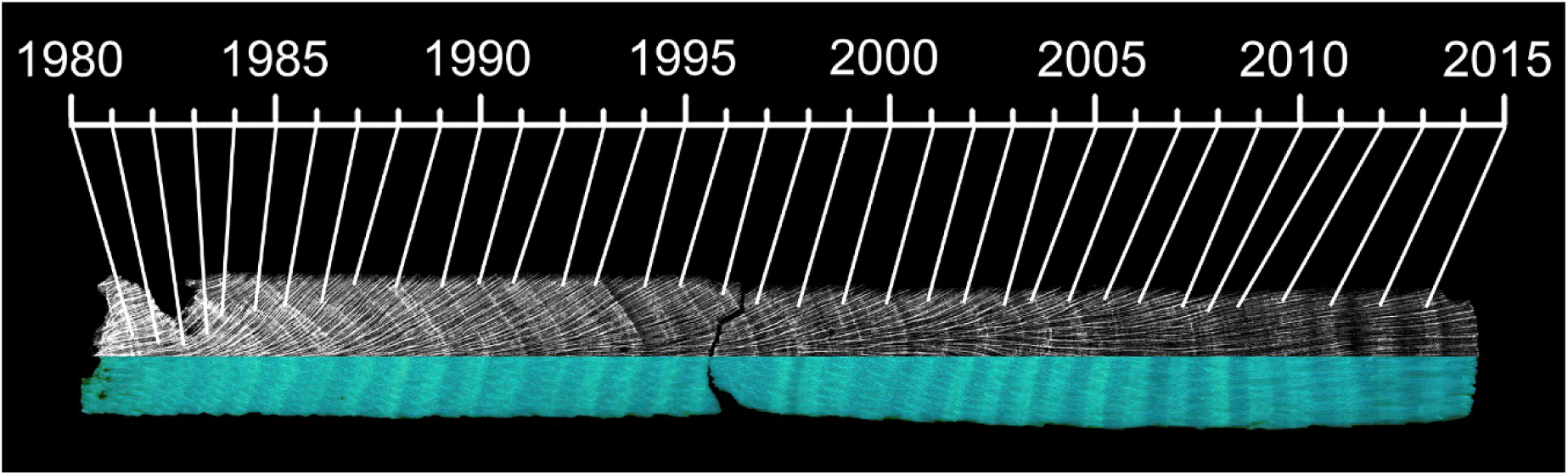
Coral growth band patterns along an *O. faveolate* core. The annual skeletal density is shown in grayscale (X-ray) as dark (low density) and light (high density) bands. Annual luminescent bands, visible under UV-A light, show bands of high and low intensity.

Since the aragonite standards used for estimating coral skeletal density in both X-ray and CT-scan methods were the same, we included X-ray derived coral density in the global coral growth analysis, assuming minimal error between both methods. The mean skeletal density of core VAR1 fell within the range of the densities measured for cores VAR2, VAR3 and VAR4 over the period of 1981-2015, suggesting that density estimates were comparable between the two methods (See Results section).

Additionally, to explore the potential relationship between growth parameters and the skeletal microstructure of the corals, measurements were taken on the exothecal dissepiments randomly along the coral slabs, including the number of annual dissepiments, their thickness, and spacing (Dodge et al. 1992; Dávalos-Dehullu et al. 2008). While the continuous measurement of the number of dissepiments per year was challenging due to changes in the trajectory of the exothecas, forty measurements of dissepiment thickness and spacing were taken for each coral specimen. The skeletal microstructure measurements were performed on digitalized images of the cores.

### Skeletal luminescence (green/blue ratio)

All coral cores were cut into slabs of 6 mm thick and treated with 13% reagent-grade NaOCl for 24 h (Nagtegaal et al. 2012). After treatment, the slabs were rinsed several times with distilled water and oven-dried at 60°C for 24 h. Annual luminescence patterns on the coral cores were visualized and photographed under UV-A light (Opalux blacklight bulb T8 20W) in a black box (Fig. 2). The red, green and blue (RGB) values were extracted from 5 mm wide tracks (62 pixels) along the coral growth axis for each digital photograph using ImageJ 1.51. The green/blue (G/B) ratio was then used as a measure of luminescence intensity in coral skeleton (hereafter skeletal luminescence). The G/B ratio reflects the influence of humic acid/aragonite ratios within the coral skeleton (Grove et al. 2010), with higher G/B intensities indicating higher freshwater discharge and sediment load (Tanzil et al. 2016). Consequently, the highest and lowest annual luminescence peaks were aligned with the months of highest (late December) and lowest (late March) water flow records, respectively (Fig. S2).

### Hydrological and environmental data

Monthly water flow data (1981‒2015) from the Canal del Dique at the Santa Helena station (ID 29037450) (Fig. 1b) were obtained from the Instituto de Hidrología, Metereología y Estudios Ambientales (IDEAM). Sea surface temperature data were not available at the temporal and spatial scales required; therefore, air temperature was used. Monthly air temperature data (1954‒2013) were obtained from the IDEAM’s Escuela Naval station (ID 14015030) (Fig. 1b). To complete the time series, data for the period 2014-2016 were extrapolated by modeling air temperature using data from the Rafael Nuñez Airport station (ID 14015080), located 6 km north of the Escuela Naval station.

The Southern Oscillation Index (SOI) data were obtained from the National Oceanic and Atmospheric Administration (NOAA) (https://www.ncdc.noaa.gov/teleconnections/enso/indicators/soi/). Negative and positive values of this index indicate El Niño and La Niña events, respectively. The Atlantic Multidecadal Oscillation (AMO) data were sourced from the Kaplan SST dataset (Enfield et al. 2001). The AMO is an index of North Atlantic temperatures that influences the growth of *O. faveolata* (Lizcano-Sandoval et al. 2019). Both SOI and AMO data were downloaded for the period 1954–2015, corresponding to the period for which the temperature data was available.

### Data treatment and analyses

Time series were obtained for skeletal density, linear extension, calcification, and luminescence. Mean values for each coral growth parameter (i.e., master chronologies) were calculated from at least two measurements per year (i.e., from two cores). Person correlations were performed between skeletal density, linear extension, and calcification rates. Linear regressions were used to detect trends over the time series of coral growth parameters, skeletal luminescence, and environmental data. Change point detection analysis was conducted to identify statistical changes between segments or states within the time series (Aminikhanghahi and Cook 2017; Truong et al. 2020). We employed the PELT algorithm for point change detection, which has been demonstrated to be efficient and accurate compared to other methods (Killick et al. 2012). Additionally, we compared the mean values of coral growth parameters and skeletal luminescence between periods before and after the major dredging events that began in 1981, using a two-tailed t-test, assuming unequal variances and an alpha level of 0.05.

The effects of environmental variables on coral growth and skeletal luminescence were tested for two periods: 1954–2015 and 1981–2015. The period 1981–2015 included air temperature, AMO, SOI, and water flow data. Water flow data was not available for the period 1954–2015. A model selection procedure was conducted to identify the best-performing predictor variables for use in a multiple linear regression, using the Akaike Information Criterion with a stepwise selection approach (Table S1; Table S3) (Quinn and Keough 2002). If none of the predictor variables outperformed a null model, the analysis was omitted (e.g., with calcification in Table S2). However, the model selection did not identify more than one predictor variable to justify running a multiple linear regression (Table S2; Table S4). As a result, simple linear regressions were performed. The response variables were centered, and predictors transformed into z-scores to standardize the data before performing the regressions, resulting in intercepts of zero. The model selection was conducted using the *Fathom* package in Matlab vR2022a, while plots and additional analyses were completed using Python. The point change detection analysis utilized the *ruptures* Python package (Truong et al. 2020).

## Results

### Coral growth parameters

The mean and standard deviation values of the skeletal density, linear extension and calcification were 0.72 ± 0.07 g cm^-3^, 1.03 ± 0.13 cm yr^-1^, and 0.74 ± 0.10 g cm^-2^ yr^-1^, respectively, corresponding to the period 1951–2015 (Table 1). The linear extension was negatively correlated to skeletal density (r = -0.29; p = 0.02; Pearson) and positively correlated to calcification rates (r = 0.69; p < 0.01; Pearson). The skeletal density was positively correlated to calcification (r = 0.46, p < 0.01; Pearson). The mean number of dissepiments per year along the coral slabs was 21.3 ± 1.3, while dissepiment thickness and spacing were 0.13 ± 0.01 mm and 0.57 ± 0.06 mm, respectively. No signs of mortality were observed along the coral cores.

**Table 1.** Coral growth mean and standard deviations from four cores of the coral *O. faveolata* in Varadero Reef.

| Core | Time span | n | Density<br>(g cm<sup>-3</sup>) | Extension<br>(cm yr<sup>-1</sup>) | Calcification<br>(g cm<sup>-2</sup> yr<sup>-1</sup>) |
| --- | --- | --- | --- | --- | --- |
| VAR1 | 1982–2015 | 34 | $0.69 \pm 0.08$ | $1.33 \pm 0.35$ | $0.91 \pm 0.24$ |
| VAR2 | 1951–2015 | 65 | $0.73 \pm 0.14$ | $1.04 \pm 0.20$ | $0.77 \pm 0.22$ |
| VAR3 | 1942–2015 | 74 | $0.80 \pm 0.12$ | $0.91 \pm 0.22$ | $0.74 \pm 0.23$ |
| VAR4 | 1950–2015 | 66 | $0.67 \pm 0.13$ | $1.02 \pm 0.20$ | $0.67 \pm 0.15$ |
| Mean | 1951–2015 | 65 | $0.73 \pm 0.07$ | $1.02 \pm 0.13$ | $0.74 \pm 0.10$ |

### Temporal trends

Skeletal density, calcification rate, and skeletal luminescence showed significant declines between 1951 and 2015 (Table 2; Fig. 3). Significant increases in water flow from 1981 to 2015 and in the AMO from 1951 to 2015 were detected (Table 2; Fig. 3). Point changes detected in 1982 for skeletal density; in 1966 and 1986 for linear extension; in 1961 for calcification rate; and in 1956, 1996, 2001, and 2011 for skeletal luminescence (Fig. 3). After 1986, linear extension exhibited greater variability among coral cores (Fig. 3). Some points of change in temperature, SOI and AMO time series occurred concurrently between 1971 and 1976 (Fig. 3), a period characterized by a low peak in air temperature and AMO, and a high peak in SOI. Both SOI and AMO showed apparent increasing trends after 1996, while air temperature data showed increases after 2001. Water flow data also depicted increases between 2006 and 2011 (Fig. 3).

**Figure 3.**
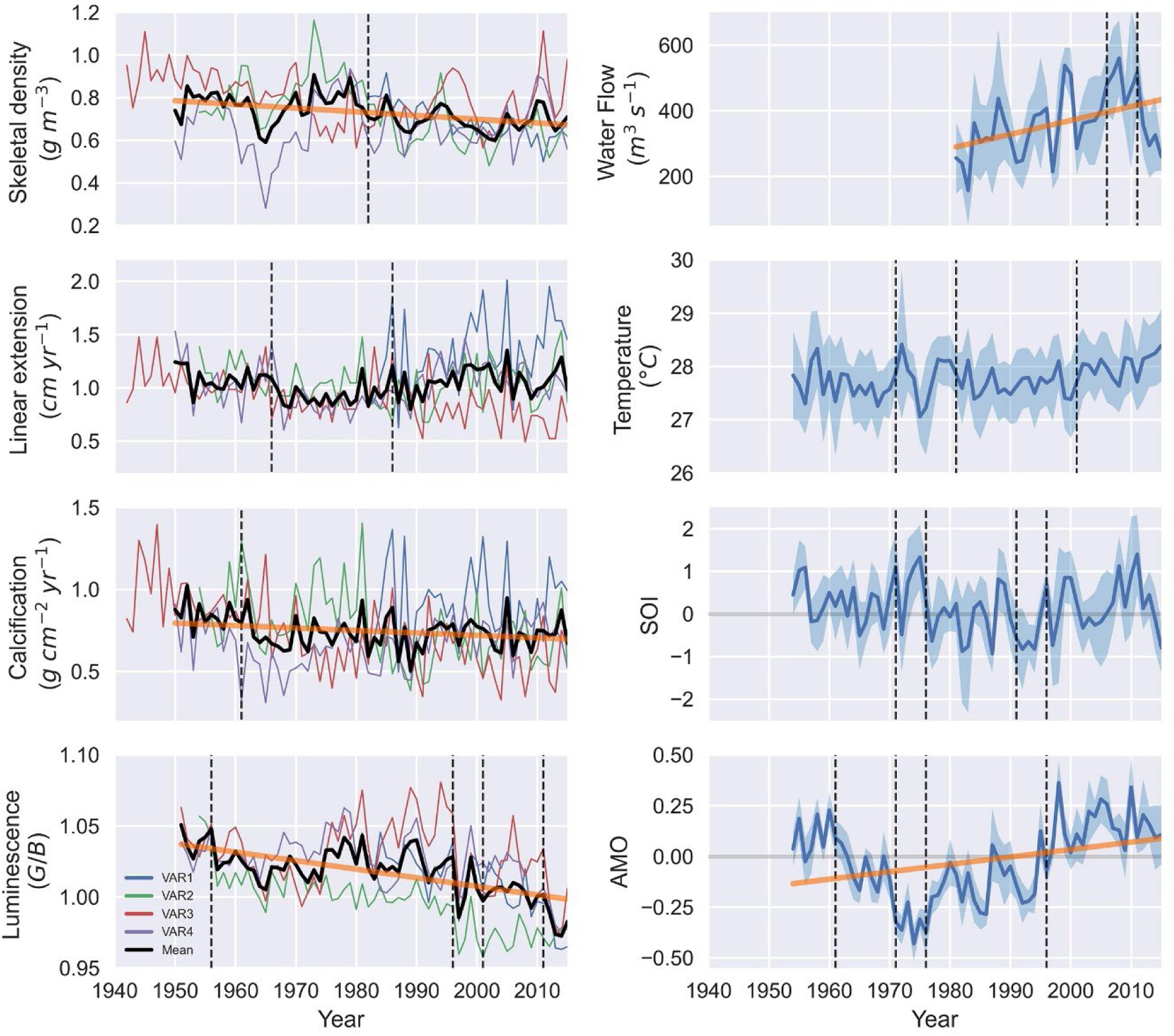
Time series of coral growth, coral luminescence, and environmental data (shaded area indicates annual standard deviations). The legend of coral cores corresponds to the left subplots. Significant trends over time are indicated by bold orange lines (linear regression: p < 0.05). Vertical dashed lines mark points of change in the time series.

**Table 2.**
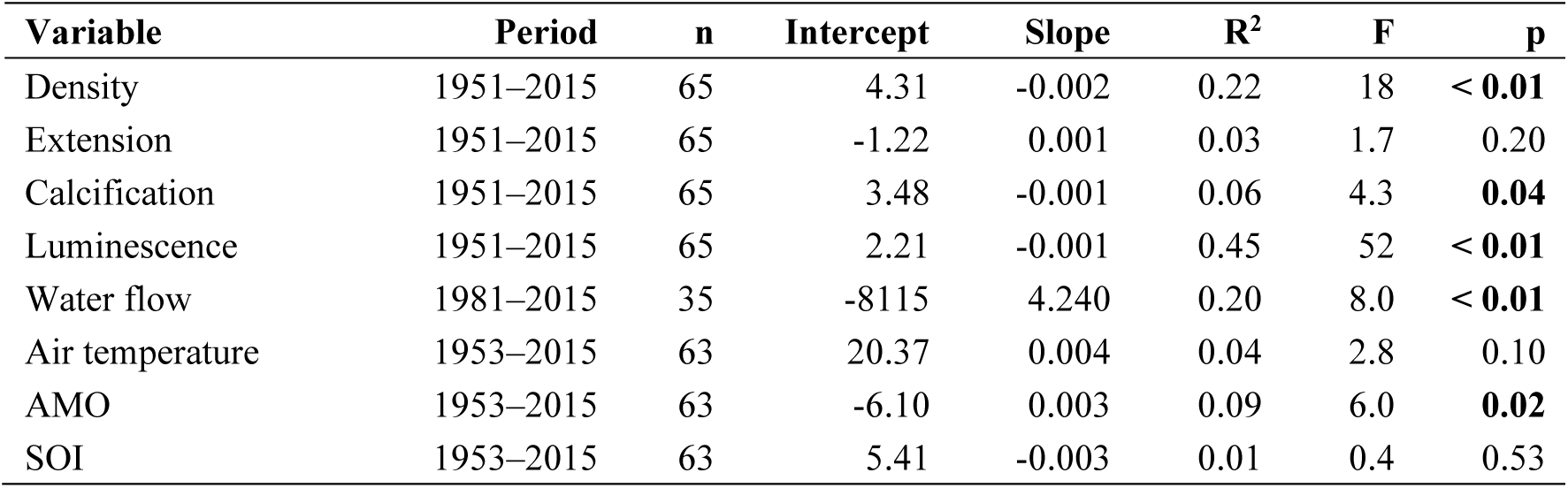
Linear regression results of mean coral growth and coral luminescence, and environmental time series data. The slopes are significantly different from zero at p < 0.05 (bold values).

| Variable | Period | n | Intercept | Slope | R<sup>2</sup> | F | p |
| --- | --- | --- | --- | --- | --- | --- | --- |
| Density | 1951–2015 | 65 | 4.31 | -0.002 | 0.22 | 18 | < **0.01** |
| Extension | 1951–2015 | 65 | -1.22 | 0.001 | 0.03 | 1.7 | 0.20 |
| Calcification | 1951–2015 | 65 | 3.48 | -0.001 | 0.06 | 4.3 | **0.04** |
| Luminescence | 1951–2015 | 65 | 2.21 | -0.001 | 0.45 | 52 | < **0.01** |
| Water flow | 1981–2015 | 35 | -8115 | 4.240 | 0.20 | 8.0 | < **0.01** |
| Air temperature | 1953–2015 | 63 | 20.37 | 0.004 | 0.04 | 2.8 | 0.10 |
| AMO | 1953–2015 | 63 | -6.10 | 0.003 | 0.09 | 6.0 | **0.02** |
| SOI | 1953–2015 | 63 | 5.41 | -0.003 | 0.01 | 0.4 | 0.53 |

The mean values for skeletal density and skeletal luminescence during the period 1951–1981 were statistically higher than those observed from 1981–2015 (t-test: p < 0.001) (Fig. 4). The mean values for linear extension were higher (t-test: p = 0.049), and calcification rates were lower (t-test: p = 0.131), during 1981–2015 compared to the period 1951–1981 (Fig. 4). In average, skeletal density was 9.9% lower, linear extension was 6.6% higher, and skeletal luminescence 1.6% less intense during 1981–2015, in comparison to the period 1951–1981.

**Figure 4.**
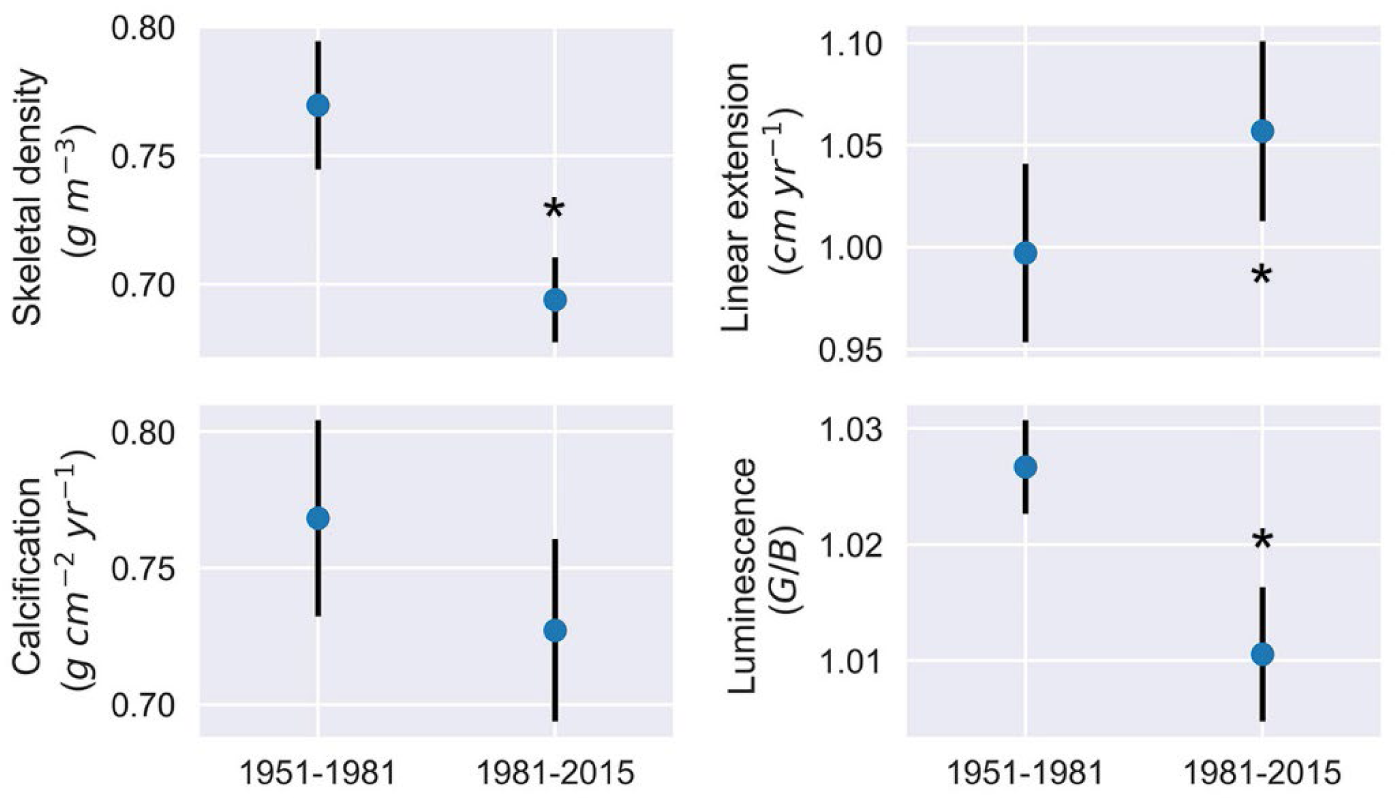
Coral growth and skeletal luminescence (mean ± CI 95%) before and after the major dredging of 1981 in the Canal del Dique. The asterisks indicate significant differences in respective variables between the two periods (t-test: p < 0.05).

### Coral responses to environmental variables

The model selection procedure indicated that predictor variables such as AMO, SOI, and air temperature were the most adequate for linear regression analysis in both periods 1954–2015 and 1981–2015 (Table S2; Table S4). Water flow data did not show significant contributions to any coral growth parameter and luminescence (e.g., R^2^ ˂ 0.03) and was not selected for linear regression for period 1981-2015 (Table S3; Table S4). The AMO explained 8% of the decrease and 19% of the increase in skeletal density and linear extension over 1954–2015, respectively (Table 3). Air temperature explained 22% of the decrease in luminescence over the same period. Over 1981–2015, the temperature significantly explained 45% of the decrease in luminescence (Table 4).

**Table 3.** Linear regression results for annual coral growth and luminescence variables against environmental predictors for the period 1954–2015. All regressions have intercept set to zero. Significant slopes are identified with a p-value of < 0.05 (bold values)

| Variable/Predictor | Slope | R<sup>2</sup> | p-value |
| --- | --- | --- | --- |
|  | *Density* |  |  |
| AMO | -0.11 | 0.08 | 0.025 |
|  | *Extension* |  |  |
| AMO | 0.30 | 0.19 | **0.001** |
|  | *Calcification* |  |  |
| SOI | 0.04 | 0.06 | 0.069 |
|  | *Luminescence* |  |  |
| Air temperature | -0.03 | 0.22 | **0.001** |

**Table 4.** Linear regression results for annual coral growth and luminescence variables against environmental predictors for the period 1981–2015. . All regressions have intercept set to zero. Significant slopes are identified with a p-value of < 0.05 (bold values)

| Variable/Predictor | Slope | R<sup>2</sup> | p-value |
| --- | --- | --- | --- |
|  | *Density* |  |  |
| AMO | -0.32 | 0.10 | 0.061 |
|  | *Extension* |  |  |
| AMO | 0.26 | 0.07 | 0.144 |
|  | *Luminescence* |  |  |
| Air temperature | -0.67 | 0.45 | **0.001** |

## Discussion

### Coral growth characteristics

The coral colonies of *O. faveolata* in Varadero Reef exhibit low skeletal density and calcification rates, while their linear extension remains relatively stable over time. The average skeletal density and calcification rates observed are the lowest reported for this species at 4 m depth, despite maintaining relatively high linear extension rates (Table 5). These corals grow under low-light conditions caused by high water turbidity, resulting in plate-like colony shapes typical of those found in light-limited environments (Pizarro et al. 2017; López-Londoño et al. 2021). The light availability and physiology of *O. faveolata* at 4 m depth in Varadero Reef are similar to those observed in *O. faveolata* colonies at 12 m depth in reefs of the Rosario Islands (Colombian Caribbean), where the quality of the optical properties of water is better (López-Londoño et al. 2021). In response to decreasing light availability with increasing depth, orbicellid corals typically increase skeletal density while reducing linear extension and calcification rates (Baker and Weber 1975; Dustan 1975; Bosscher 1993). However, the skeletal growth characteristics of *O. faveolata* in Varadero Reef deviate from expected patterns for corals growing under low-light conditions. Correlations between the growth variables suggest that the observed growth strategy of *O. faveolata* focuses on utilizing calcification products to achieve faster vertical growth with less dense skeletons, contrary to previous findings for orbicellid corals (Carricart-Ganivet and Merino 2001). The benefits of faster growth in the shallow suboptimal conditions of Varadero Reef (∼2–6 m depth for *O. faveolata*) require further research.

**Table 5.** Linear extension, skeletal density, and calcification rates of *O. faveolata* over relatively long periods and different depths in the Caribbean Sea.

| Location | Period | Depth (m) | Number of cores | Extension (cm yr<sup>-1</sup>) | Density (g cm<sup>-3</sup>) | Calcification (g cm<sup>-2</sup> yr<sup>-1</sup>) | Reference |
| --- | --- | --- | --- | --- | --- | --- | --- |
| Veracruz Reef, Mexico | 1835–2002 |  | 1 | 1.12 ± 0.23 |  |  | (Horta-Puga and Carriquiry 2014) |
| Upper Florida Keys, USA | 1937–1996 |  | 7 | 0.79 ± 0.07 | 1.18 ± 0.06 | 0.91 ± 0.06 | (Helmle et al. 2011) |
| Turneffe Islands, Belize | 1954–1996 | 2 | 1 | 0.88 ± 0.07 | 0.98 ± 0.05 | 0.83 ± 0.11 | (Zoppe and Gischler 2024) |
| Glovers Reef, Belize | 1899–2005 | 4 | 1 | 0.95 ± 0.09 | 1.13 ± 0.06 | 1.07 ± 0.11 | Zoppe and Gischler (2024) |
| **Varadero Reef, Colombia** | **1951–2015** | **4** | **4** | **1.02 ± 0.13** | **0.73 ± 0.07** | **0.74 ± 0.10** | **This study** |
| Cheeca Rocks Reef, USA | 2004–2013 | 4–5 | 11 | 0.89 ± 0.07 | 1.25 ± 0.03 | 1.10 ± 0.08 | (Manzello et al. 2015) |
| Barrier reef, Belize | 1942–1999 | 4.8 | 1 | 1.20 ± 0.01 | 0.99 ± 0.05 | 1.19 ± 0.10 | Zoppe and Gischler (2024) |
| Little Conch Reef, USA | 2004–2013 | 5–6 | 12 | 0.73 ± 0.03 | 1.34 ± 0.01 | 0.97 ± 0.03 | (Manzello et al. 2015) |
| Serrana Cay, Colombia | 1963–2015 | 4–9 | 5 | 0.96 ± 0.11 | 1.08 ± 0.08 | 1.02 ± 0.12 | (Lizcano-Sandoval et al. 2019) |
| Cayo Arenas, Mexico | 1992–2016 | 7–12 | 5 | 0.82 ± 0.08 | 1.04 ± 0.30 | 0.85 ± 0.18 | (Sánchez-Pelcastre et al. 2023) |
| Flower Garden Banks, USA | 1970–2014 | 19 | 10 | 0.71 ± 0.04 | 1.42 ± 0.03 | 0.99 ± 0.05 | (Manzello et al. 2021) |

### Skeletal microstructure characteristics

The mean number of dissepiments per year (21.3 ± 1.3) was nearly double that observed in other studies. *O. faveolata* specimens from Mahahual, Mexico and La Parguera, Puerto Rico recorded annual averages of 13.6 ± 0.9 (Dávalos-Dehullu et al. 2008) and 11.8 ± 1.6 (Winter and Sammarco 2010) exothecal dissepiments, respectively. The dissepiment spacing observed in this study (0.57 ± 0.06 mm) was similar to that found by Dávalos-Dehullu et al. (2008), but the dissepiments in Varadero were almost twice as thick (0.13 ± 0.01 mm). The high number of dissepiments formed annually could explain the low skeletal density observed. The decline in skeletal luminescence intensity after 1981 may be related to the decrease in skeletal density, as skeletal microstructure contributes a small proportion to luminescence (Barnes and Taylor 2005). However, the causes of variations in skeletal microstructure remain unclear. Some studies suggest that these variations are due to seasonal changes in tissue thickness and calcification rates, which affect coral density banding patterns (Barnes and Lough 1993; Cruz-Piñón et al. 2003; Dávalos-Dehullu et al. 2008).

Lunar cycles have been proposed as a trigger for dissepiment formation, with one dissepiment forming per full moon on average (∼12 annually) (Dávalos-Dehullu et al. 2008; Winter and Sammarco 2010). Corals can respond to lunar cycles by detecting the blue portion of lunar irradiance (Gorbunov and Falkowski 2002), but in Varadero Reef, the optical properties of the water are variable and suboptimal, particularly in the blue light spectrum (López-Londoño et al. 2021). If irradiance at 4 m depth in Varadero Reef represent between 20% and 30% of surface irradiance (Pizarro et al. 2017; López-Londoño et al. 2021), then moon light irradiance may range between 6 x 10^-4^ and 9 x 10^-4^ µmol quanta m^-2^ s^-1^, assuming zenith moon light irradiance at the surface is ∼3 x 10^-3^ µmol quanta m^-2^ s^-1^ (Winter and Sammarco 2010). This hypothetical moon light irradiance at 4 m depth is below the coral photoreception sensitivity threshold of ∼2 x 10^-3^ µmol quanta m^-2^ s^-1^ (Gorbunov and Falkowski 2002). However, daily variations in water quality can cause large fluctuations in light attenuation coefficients (Kd) in Varadero Reef (López-Londoño et al. 2021), which may temporarily enable corals to respond to light. The causes of variation in skeletal microstructure for *O. faveolata* in Varadero Reef are likely multifaceted. Further investigation into the effects of light on coral skeletal microstructure is needed.

### Growth responses to environmental variables

The AMO significatively accounted for 19% of the increases in linear extension over the period 1954-2015. The growth of *O. faveolata* has shown to be related to the AMO to some extent, though the patterns may vary. Our results are comparable to those from the remote reefs of Serrana Cay, Colombian Caribbean, where AMO showed long-term influence in the skeletal growth of *O. faveolata* (Lizcano-Sandoval et al. 2019). However, contrasting patterns were found in *O. faveolata* corals from Belize and Florida, where linear extension and skeletal density were negatively and positively correlated to the AMO, respectively (Helmle et al. 2011; Zoppe and Gischler 2024). Air temperature explained 22% of the decreases in skeletal luminescence in 1954-2015 and 45% in the period 1981-2015.

No significant contribution was observed between changes in water flow and coral growth or luminescence variables after 1981 (Table S3), yet changes in the time series trends of skeletal density and linear extension were detected in 1982 and 1986, respectively (Fig. 3). The mean values of skeletal density, linear extension, and luminescence in the period 1981–2015 were significantly different from those of the period 1951–1981 (Fig. 4), suggesting a likely response of *O. faveolata* to increased water flow and decreased water quality due to dredging activities from 1981 to 1984, which removed 18.8 million m^3^ of soil and sediments (Mogollón 2013). The points of change detected in the skeletal luminescence time series are apparently related to El Niño from 1997–1998 and La Niña from 1998–2001, which coincided with low and high water flow during those periods, respectively (Fig. 3). The air temperature data showed a decrease that coincided with La Niña from 1973–1976, together with slight increases in skeletal density and luminescence, and decreases in linear extension.

The corals in Varadero Reef are exposed to multiple stressors that can affect their skeletal density, linear extension, and calcification rates. Limited photosynthetic potential, combined with the presence of organic matter in the water column, can cause a shift towards increased heterotrophy to meet energetic requirements. For some tropical corals, heterotrophy can contribute 15% to 35% of daily energy requirements (Houlbrèque and Ferrier-Pagès 2009). Under heterotrophic conditions, linear extension may double, skeletal density can decrease, and skeletal mass may increase (Ferrier-Pagès et al. 2003; Fabricius 2005). Cartagena Bay also experiences seasonal hypoxic conditions, with low dissolved oxygen levels (< 4 mg L^-1^) occurring between 10 and 20 m depth, but during transitional seasons (May-August), these levels can be found at 5-10 m depth (Tosic et al. 2019). Prolonged exposure to hypoxia can lead to coral bleaching, tissue loss, or reduction in calcification rates (Pezner et al. 2023). Additionally, heavy metal concentrations (e.g., As, Zn, Cu, Ni, Cr, Mn, Fe, Pb) in Cartagena Bay often exceed acceptable standards (Restrepo et al. 2016; Tosic et al. 2019). Concentrations of total suspended solids (6-72 mg L^-1^), nitrates (5-389 µg L^-1^), and phosphates (17-140 µg L^-1^) can also surpass thresholds considered healthy for coral ecosystems (Fabricius 2005; Tosic et al. 2019). A potential threat for coral growth in Varadero Reef that may require more attention is sea-level rise. Cartagena presents one of the largest sea-level rise rates in the Caribbean, which have accelerated between 2000 and 2019 with a rate of 7.02 mm yr^-1^ (Restrepo-Ángel et al. 2021).

The absence of signs of mortality in any of the coral cores raise the question of how these corals have survived and apparently thrived for over 65 years under the influence of the Canal del Dique. However, the constant increase in water flow suggests a decline in water quality, which poses an ongoing threat to coral health and survival in Varadero Reef. The reason for an accelerated linear extension might be a response to the particular conditions of light and pollution, but further research is required to understand coral growth and responses to these conditions in Cartagena Bay. A more frequent and comprehensive water quality monitoring program is needed to assess the effects of the multiple stressors on coral growth, and conservation efforts should be implemented in the bay to protect Varadero Reef.

