## Supplemental Information for "Accelerated growth of the scleractinian coral *Orbicella faveolata* under turbid and polluted water conditions"

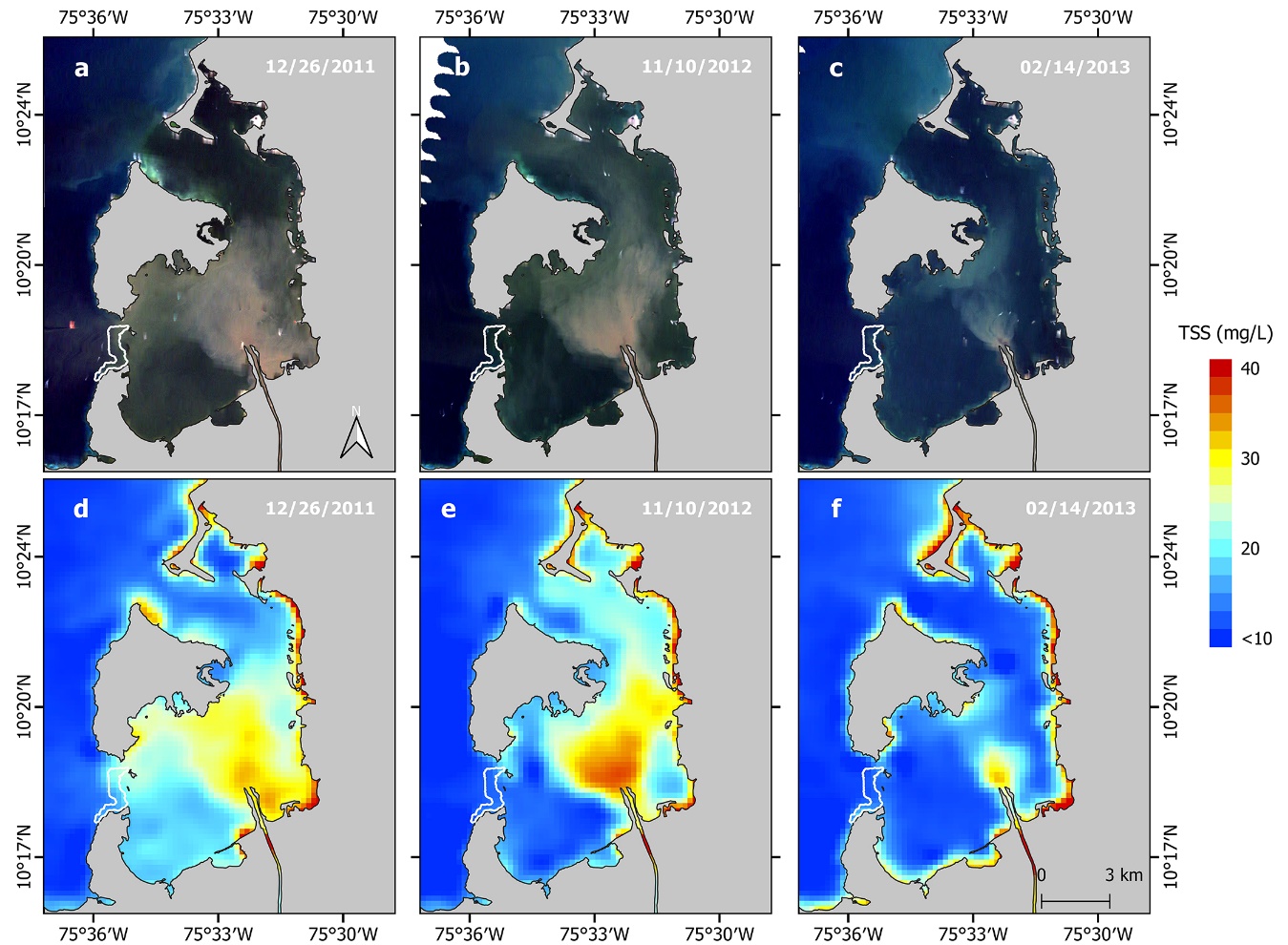

**Figure S1.** Total suspended sediments (TSS) within Cartagena Bay according to three water flow rates recorded upstream the Canal del Dique (Santa Helena Station): high (632 m^3^ s^-1^) on December 26, 2011, moderate (378 m^3^ s^-1^) on November 10, 2012, and low (158 m^3^ s^-1^) on February 14, 2013. True-color satellite imagery showing different levels of suspended sediments in Cartagena Bay (a-c). MODIS Terra (MOD09GQ; band 1) satellite imagery calibrated for TSS following the model by Restrepo et al. (2016) (d-f). The Varadero coral reef is outlined by a white-bordered polygon.

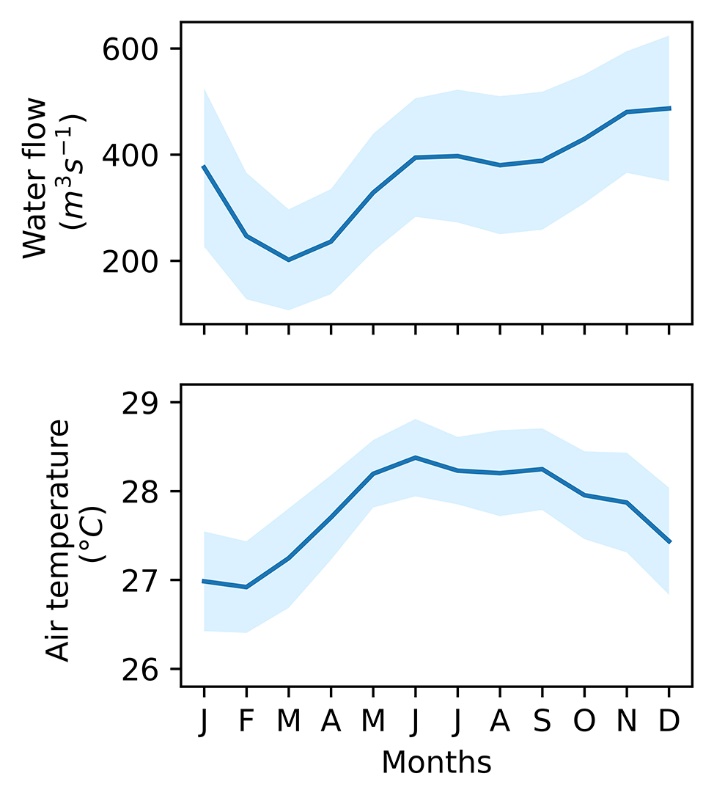

**Figure S2.** Climatology of water flow (1981-2015) and air temperature (1954-2015) monthly data from the Canal del Dique and Cartagena Bay (1950-2015). The data sources are detailed in the methods section.

The model selection is a procedure that first runs marginal (independent) tests on each variable, including a test with no variables (null model). The best variables are selected based on the Akaike Information Criterion (AIC), weights and ratio values. The weight indicates the probability that the respective variable is the best one to be included in a model. The ratios compare each variable against the best (first) variable, indicating how many times it is better than the others. Then, conditional tests are performed with the variables that indicated to be better than the null model. If a null model is better than any other model then the procedure is stopped (e.g., no conditional tests are required for calcification after the results in Table S3). The conditional tests are the last step to select the predictor variables to include in the optimized multiple linear regression analysis.

**Table S1.** Results of the marginal tests (independent) with AIC on environmental predictors for coral growth parameters and luminescence (period 1954-2015). Sum of squares residuals (RSS) are additive.

| **Variable** | **RSS** | **R^2^** | **AIC** | **ΔAIC** | **Weights** | **Ratio** |
| --- | --- | --- | --- | --- | --- | --- |
| *Density* | | | | | | |
| AMO | 0.29 | 0.08 | -329.1 | 0.0 | 0.7 | 1.0 |
| SOI | 0.30 | 0.04 | -326.2 | 2.9 | 0.2 | 4.3 |
| *None* | 0.31 |  | -326.1 | 3.0 | 0.1 | 4.5 |
| Temperature | 0.31 | 0.00 | -324.0 | 5.1 | 0.1 | 12.8 |
| *Extension* | | | | | | |
| AMO | 0.84 | 0.19 | -262.8 | 0.0 | 1.0 | 1.0 |
| *None* | 1.03 |  | -252.2 | 10.6 | 0.0 | 204 |
| SOI | 1.02 | 0.01 | -250.6 | 12.3 | 0.0 | 462 |
| Temperature | 1.02 | 0.01 | -250.5 | 12.3 | 0.0 | 465 |
| *Calcification* | | | | | | |
| SOI | 0.55 | 0.06 | -288.8 | 0.0 | 0.6 | 1.0 |
| *None* | 0.58 |  | -287.1 | 1.7 | 0.2 | 2.3 |
| AMO | 0.58 | 0.01 | -285.8 | 3.0 | 0.1 | 4.5 |
| Temperature | 0.58 | 0.00 | -285.0 | 3.8 | 0.1 | 6.8 |
| *Luminescence* | | | | | | |
| Temperature | 0.01 | 0.22 | -520.5 | 0.0 | 1.0 | 1.0 |
| AMO | 0.01 | 0.11 | -512.3 | 8.2 | 0.0 | 61 |
| SOI | 0.02 | 0.04 | -507.2 | 13.3 | 0.0 | 769 |
| *None* | 0.02 |  | -507.0 | 13.5 | 0.0 | 840 |

**Table S2.** Results of the conditional tests with AIC on environmental predictors for coral growth parameters and luminescence (period 1954-2015). Sum of squares residuals (RSS) are additive.

| **Variable** | **RSS** | **R^2^** | **AIC** | **ΔAIC** | **Weights** |
| --- | --- | --- | --- | --- | --- |
| *Density* | | | | | |
| AMO | 0.29 | 0.08 | -329.1 | 3.0 | 0.7 |
| None | 0.29 |  | -329.1 | 0.5 | 0.3 |
| *Extension* | | | | | |
| AMO | 0.84 | 0.19 | -262.8 | 10.6 | 1.0 |
| None | 0.84 |  | -262.8 | 0.0 | 0.5 |
| *Calcification* | | | | | |
| SOI | 0.55 | 0.06 | -288.8 | 1.7 | 0.6 |
| None | 0.55 |  | -288.8 | 0.0 | 0.4 |
| *Luminescence* | | | | | |
| Temperature | 0.01 | 0.22 | -520.5 | 13.5 | 1.0 |
| None | 0.01 |  | -520.5 | 0.2 | 0.4 |

**Table S3.** Results of the marginal tests (independent) with AIC on environmental predictors for coral growth parameters and luminescence (period 1981-2015). Sum of squares residuals (RSS) are additive.

| **Variable** | **RSS** | **R^2^** | **AIC** | **ΔAIC** | **Weights** | **Ratio** |
| --- | --- | --- | --- | --- | --- | --- |
| *Density* | | | | | | |
| AMO | 30.5 | 0.10 | -0.4 | 0.0 | 0.5 | 1.0 |
| *None* | 34.0 |  | 1.1 | 1.5 | 0.2 | 2.1 |
| Temperature | 33.6 | 0.01 | 3.0 | 3.4 | 0.1 | 5.4 |
| Water Flow | 33.9 | 0.00 | 3.2 | 3.6 | 0.1 | 6.1 |
| SOI | 34.0 | 0.00 | 3.4 | 3.7 | 0.1 | 6.5 |
| *Extension* | | | | | | |
| AMO | 31.7 | 0.07 | 0.9 | 0.0 | 0.3 | 1.0 |
| *None* | 34.0 |  | 1.1 | 0.2 | 0.3 | 1.1 |
| SOI | 32.4 | 0.05 | 1.7 | 0.8 | 0.2 | 1.5 |
| Water Flow | 32.8 | 0.03 | 2.1 | 1.2 | 0.2 | 1.9 |
| Temperature | 33.2 | 0.02 | 2.5 | 1.6 | 0.1 | 2.3 |
| *Calcification* | | | | | | |
| *None* | 34.0 |  | 3.0 | 1.9 | 0.4 | 1.0 |
| SOI | 32.6 | 0.04 | 3.4 | 2.3 | 0.3 | 1.5 |
| Water Flow | 33.7 | 0.01 | 1.9 | 0.8 | 0.1 | 2.6 |
| AMO | 34.0 | 0.00 | 3.3 | 2.2 | 0.1 | 3.0 |
| Temperature | 34.0 | 0.00 | 1.1 | 0.0 | 0.1 | 3.1 |
| *Luminescence* | | | | | | |
| Temperature | 18.59 | 0.45 | -17.8 | 0.0 | 1.0 | 1.0 |
| AMO | 25.43 | 0.25 | -6.8 | 11.0 | 0.0 | 241.4 |
| *None* | 34.00 |  | 1.1 | 18.9 | 0.0 | 12609.0 |
| SOI | 33.86 | 0.00 | 3.2 | 21.0 | 0.0 | 36153.0 |
| Water Flow | 33.96 | 0.00 | 3.3 | 21.1 | 0.0 | 38081.0 |

**Table S4.** Results of the conditional tests with AIC on environmental predictors for coral growth parameters and luminescence (period 1981-2015). Sum of squares residuals (RSS) are additive. Tests for calcification were skipped according to marginal tests results.

| **Variable** | **RSS** | **R^2^** | **AIC** | **ΔAIC** | **Weights** |
| --- | --- | --- | --- | --- | --- |
| *Density* | | | | | |
| AMO | 0.1 | 0.07 | 0.5 | 1.5 | 0.5 |
| None |  |  | 0.4 | 0.0 | 0.4 |
| *Extension* | | | | | |
| AMO | 31.7 | 0.07 | 0.9 | 0.3 | 0.2 |
| None | 31.7 |  | 0.9 | 0.3 | 0.0 |
| *Luminescence* | | | | | |
| Temperature | 18.6 | 0.45 | -17.8 | 1.0 | 18.9 |
| None | 18.6 |  | -17.8 | 0.4 | 0.0 |
